# Correcting for Dependent P-values in Rhythm Detection

**DOI:** 10.1101/118547

**Authors:** Alan L. Hutchison, Aaron R. Dinner

## Abstract

There is much interest in using genome-wide expression time series to identify circadian genes. Several methods have been developed to test for rhythmicity in sparsely sampled time series typical of such measurements. Because these methods are statistical in nature, they rely on estimating the probabilities that patterns arise by chance (i.e., p-values). Here we show that leading methods implicitly make inappropriate assumptions of independence when estimating p-values. We show how to correct for the dependence to obtain accurate estimates for statistical significance during rhythm detection.

## 1 Introduction

Molecular rhythms are now commonly identified by statistically analyzing whole-genome expression time series measurements. To this end, a number of rhythm detection algorithms have been introduced [14, 6, 15, 13, 3, 16, 5, 8]. Although different algorithms are sensitive to different data features, they all seek to estimate the probabilities that perceived patterns arise by chance from experimental uncertainties (i.e., p-values). Because the p-values are often compared with desired significance thresholds to determine which genes are considered rhythmic, accurate p-value estimates are important.

A p-value is the probability that a test statistic is observed when the null hypothesis is true. For rhythm detection methods, the null hypothesis is generally that a time series consists of independent draws from a normal distribution. The null distribution corresponds to the distribution of test statistics that the algorithm yields when applied to time series generated under the null hypothesis. A p-value is the fraction of samples from this distribution that are greater than or equal to the test statistic value for the time series of interest. By construction, the *N* p-values for time series generated under the null distribution should be uniformly distributed between 0 and 1 because the *i*-th highest test statistic is larger than *N* − *i* values. Non-uniformly distributed p-values for time series generated in a fashion consistent with the null hypothesis indicate that the p-value estimates are not accurate.

Many popular rhythm detection algorithms do not yield accurate p-values. For example, in Hutchison *et al.* [6], we examined JTK_CYCLE [5]. There, we showed that testing different phases introduces a multiple-hypothesis testing problem and that the Bonferroni correction employed in the original implementation of the method [5] results in p-values that are too large. To correct for this issue, as well as the effect of selecting the best phase, we empirically sampled the null distribution. The resulting algorithm, which we termed empirical JTK (eJTK), outperforms the original algorithm on simulated data [6], demonstrating the practical utility of accurate p-values.

In this paper, we pursue this issue much further. We show that the RAIN method [13] requires empirical evaluation of the null distribution, similar to JTK_CYCLE. The fundamental issue is the same as discussed for JTK_CYCLE: tests of a time series with different reference waveforms are not independent. We then show that a related but distinct issue arises in MetaCycle [14], which combines results from multiple rhythm detection methods. There too the p-values are not independent. In that case, the Brown adjustment to Fisher Integration [2] is sufficient for obtaining accurate p-values. Interestingly, we find that, with accurate p-values, MetaCycle underperforms the individual methods on which it is based (Lomb-Scargle [9, 11], ARSER [16], and JTK_CYCLE [5]). This result suggests that combining p-values can decrease sensitivity by effectively averaging the output of individual tests.

## 2 Results

### 2.1 Comparisons of multiple reference waveforms to a single time series are not independent

JTK_CYCLE [5], RAIN [13], and eJTK [6] all use reference waveforms with different phases (and in the case of RAIN and eJTK, different peak to trough and trough to peak times) to find the reference waveform that best matches the experimental waveform according to a test statistic (Kendall's τ for JTK_CYCLE and eJTK, Mann-Whitney U for RAIN). The three methods, however, take different approaches to obtaining a single p-value for the time series from the many p-values obtained for each reference waveform comparison. JTK_CYCLE applies the Bonferroni correction, which was developed to control the Family-Wide Error Rate: the probability of obtaining one or more false positive results. The Bonferroni correction multiplies the best-matching p-value by the number of comparisons to produce an adjusted p-value that is too large, as described in Hutchison *et al.* 2015 [6].

RAIN instead uses the Benjamini-Hochberg method [1] (also known as the q-value approach [12]), which was developed to control the False Discovery Rate: the proportion of positive results that are false. As shown in the context of JTK_CYCLE in Hutchison *et al.* 2015, [6], the Benjamini-Hochberg method also results in inaccurate p-values. The reason is that the Benjamini-Hochberg method assumes that the input p-values are independent. However, the p-values from different reference waveforms are correlated since they all concern the same experimental time series. Running RAIN on simulated data generated from independent draws from the normal distribution (i.e., under the null hypothesis) results in p-values that are not uniformly distributed (Fig. 1A).

**Figure 1:**
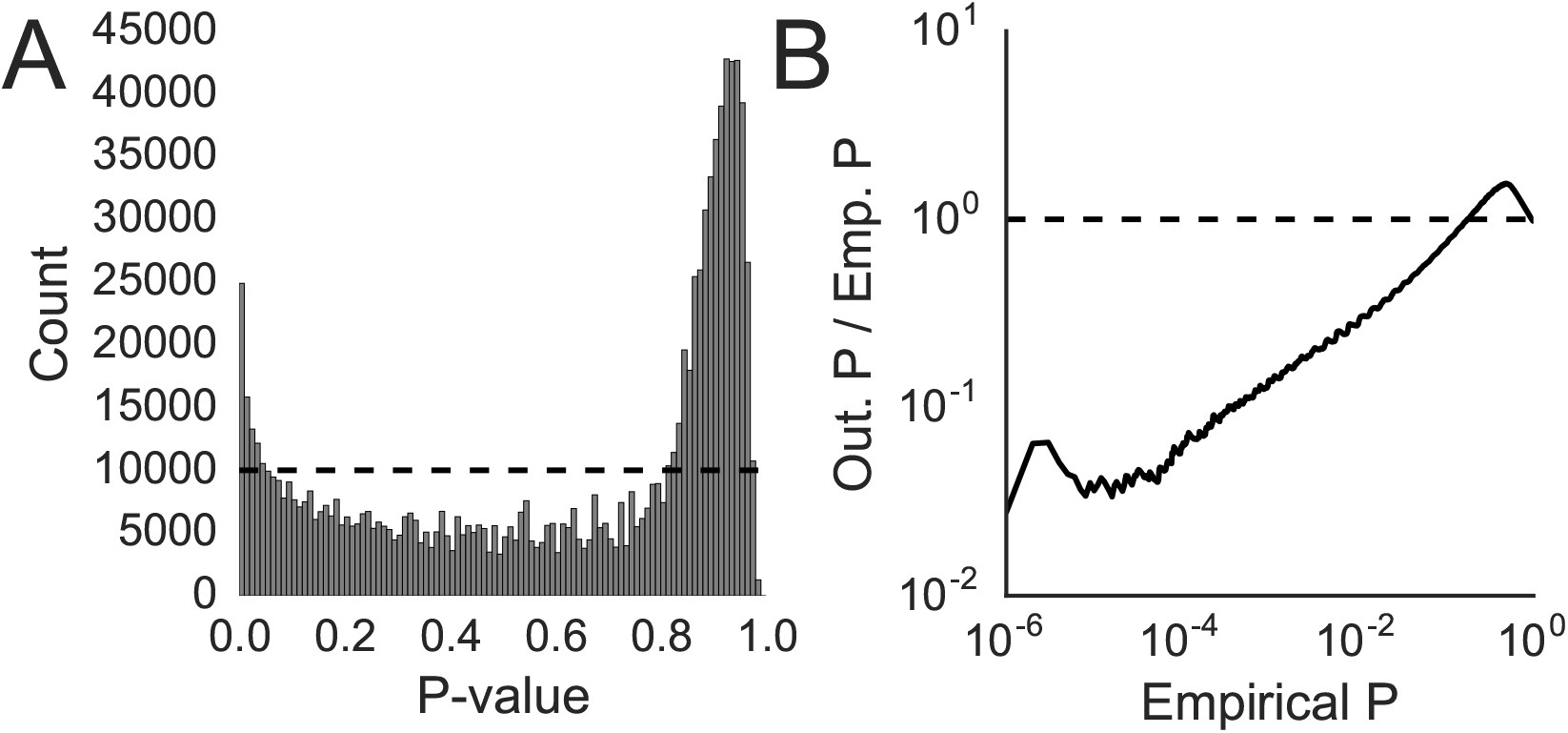
RAIN does not produce p-values that have a uniform distribution under the null hypothesis. (A) The p-values produced by RAIN run on 10^6^ Gaussian noise time series (the null distribution) are not uniform (the dashed line). (B) The ratio, *r*, of the output p-values to empirical estimates (see Eqs. 1 and 2). Each time series consisted of 24 points drawn from a Gaussian distribution, simulating data generated every 2 hours over 48 hours. RAIN was run with a time point spacing of 2 hours and a period of 24 hours.

For the reasons discussed in the Introduction, for *N* time series generated under the null distribution, the test statistic with *i* test statistics higher than it should have p-value

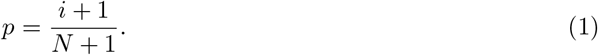

Consequently, we can compare the output p-values to empirical p-value estimates (i.e., Eq. 1) by calculating the ratio

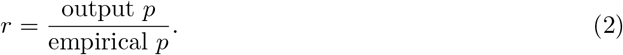

The results are shown in Fig. 1B. Ideally, a method should yield a horizontal line at *r* = 1 (the dashed line in Figure 1B). Instead, consistent with the left peak in Fig. 1A, low p-values are smaller than they should be (*r* < 1), and, consistent with the right peak in Fig. 1A, high p-values are larger than they should be (*r* > 1). The former case (*r* < 1 for low p-values) tends to produce false positives—rhythmic patterns are accorded more significance than they should be. A solution to this problem is to use the empirical p-values to judge the significance of RAIN results.

### 2.2 P-values produced by MetaCycle are not uniform under the null distribution

Given that there are a variety of rhythm detection methods available that focus on different aspects of experimental time series, it seems like it should be advantageous to combine methods. Recently, Wu *et al.* [14], implemented this idea by using Fisher integration [4] to combine the p-values from JTK_CYCLE [5], Lomb-Scargle [9, 11], and ARSER [16] in a method called MetaCycle. In this section, we analyze this method and its component algorithms from the perspective of the previous section. In the following section, we consider the Fisher integration explicitly.

We generated 1000 time series of 12 time points spaced at 2-hour intervals by drawing values from the normal distribution. We did not include replicates for time points because ARSER is only employed in MetaCycle when replicates are not available, and we wanted to test the three-algorithm combination. Consistent with Hutchison *et al.* 2015 [6], the p-values produced by JTK_CYCLE were too large. The p-values from LS were also too large (Fig. 2A), but those from ARSER were reasonably accurate (r ≈ 1). However, the p-values for the combination (Meta2D) are non-uniformly distributed, with underestimates of the p-values for low values and overestimates for high values (Fig. 2A).

**Figure 2:**
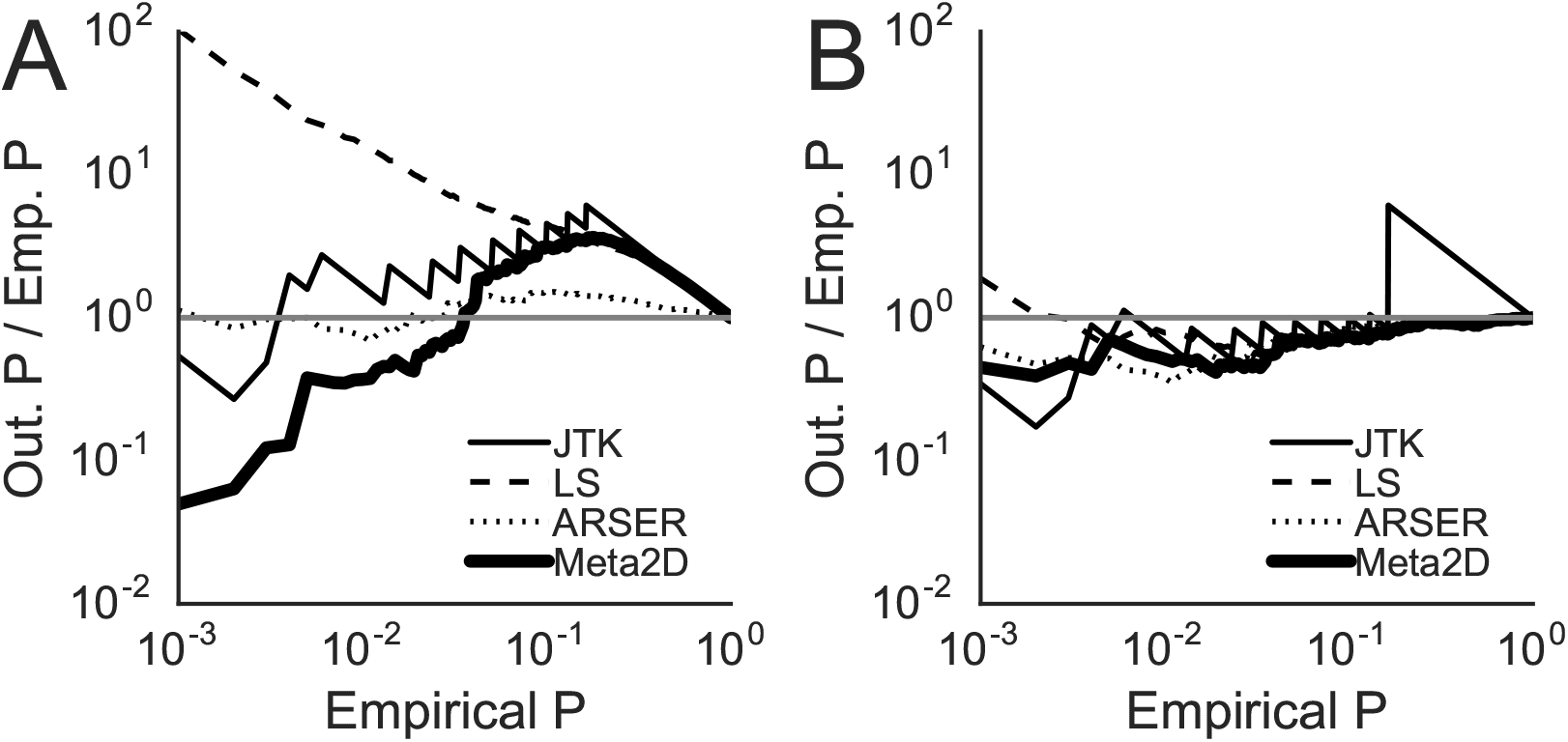
MetaCycle was run on 1000 time series generated randomly from a normal distribution, producing p-values for JTK_CYCLE (JTK), Lomb-Scargle (LS), ARSER, and their Fisher integration (Meta2D). (A) The p-values generated by the methods from MetaCycle, with the exception of those from ARSER, deviate from the uniform distribution. (B) Using the empirical method detailed in the text and in Hutchison *et al* 2015 [6], we corrected the p-values of the methods from MetaCycle such that they more closely match the uniform distribution. JTK_CYCLE sets many p-values equal to 1, leading to the deviation from 1 for high p-values as the empirical method cannot uniformly redistribute those p-values.

To make these data more accurate, we took an analogous approach to above and to Hutchison *et al* 2015 [6]. We ran MetaCycle on 1,000,000 randomly generated time series and used those results to adjust the p-values from the 1000 time series for the four different methods (LS, JTK, ARSER, and Meta2D). Although the resulting p-values are markedly improved (as evidenced by the fact that r ≈ 1 in Fig. 2B), significant deviations from the emprical values remain. In particular, for JTK_CYCLE there is a ceiling of 1 applied to the Bonferroni-corrected p-values that are output, and this results in a loss of information that cannot be fixed, giving rise to the peak at high p-values in Fig. 2B.

### 2.3 P-values from different rhythm detection methods for the same time series are not independent

Fisher integration assumes that the input p-values are independent, but the application of multiple methods to the same experimental time series yields correlated values. To illustrate this issue, we examine Fisher Integration with simulated data for two limiting cases. In the first case, we generate 3 sets (*A*, *B,* and *C*) of 1000 p-values by randomly drawing from a uniform distribution from 0 to 1 (i.e., 3 datasets generated under the null hypothesis). Drawing the points in this manner guarantees that they are uncorrelated from one another. In Fig. 3A, we combine these data with Fisher Integration to create a new set *D,* where *D_i_* (*i* = 1,…, 1000) is the combination of *A_i_*, *B_i_*, and *C_i_*. We combine the p-values and, as desired, obtain r ≈ 1.

**Figure 3:**
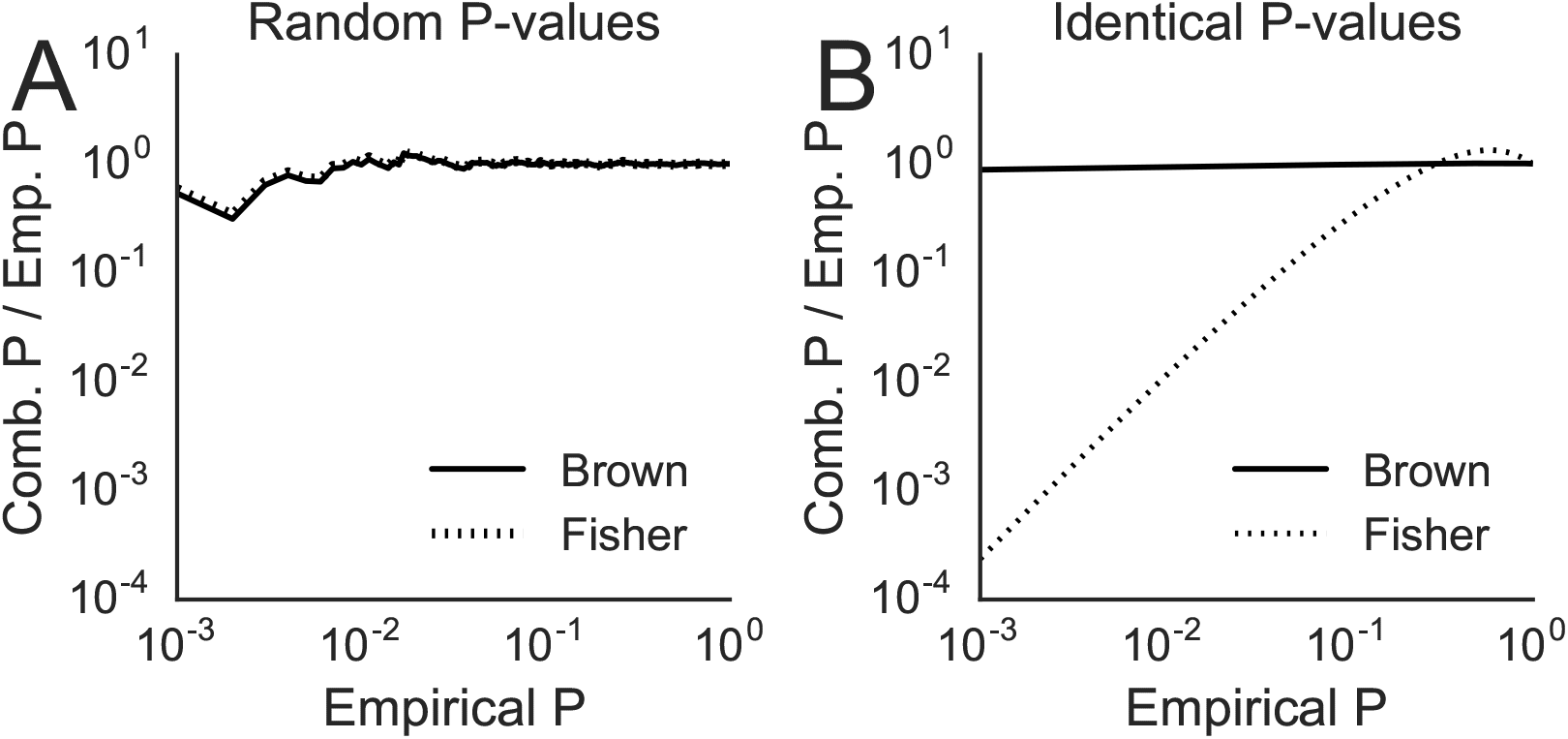
Illustration of the effects of dependent p-values during integration. (A) P-values from integration of three sets of randomly generated p-values, combined using the Fisher method (dashed line) or the Brown method (solid line). (B) P-values from the integration of three sets of identical evenly distributed p-values, combined using the Fisher method (dashed line) or the Brown method (solid line).

In the second case, we generate 3 sets of 1000 p-values that are each identical, making the correlation between the sets 1. Using the Fisher method in this case (dashed line, Fig. 3B) results in p-values with *r* ≪ 1. To correct for the correlation, we adopt the Brown method, a modification of the Fisher method for one-sided tests of significance [2]. The Brown method uses the covariance between the different p-value sets to adjust the combined test statistic (*χ*^2^) distribution and produce adjusted p-values. In the case where the p-values are independent, it provides the same results as the Fisher method (Fig. 3A). When the p-values are not independent, however, the Brown method produces accurate integrated p-values with *r* ≈ 1 (Fig. 3B), indicating that they are uniformly distributed from 0 to 1.

### 2.4 Properly combining p-values does not appear to result in improved rhythm detection

Having discussed how to adjust the methods from MetaCycle to produce accurate p-values and how to use Brown's method for accurate integration, we now explore the effects of the Fisher method and the Brown method on combining different methods. As noted above, the former leads to problems when the p-values produced by different methods are correlated. For a dataset of 12 time points drawn from a Gaussian distribution, the *R*^2^ between empirically-corrected p-values produced by JTK_CYCLE and Lomb-Scargle is *R*^2^ = 0.388, that between JTK_CYCLE and ARSER is *R*^2^ = 0.392, and that between Lomb-Scargle and ARSER is *R*^2^ = 0.999. As a result, pairwise Fisher Integration yields p-values roughly 1/100-th to 1/1000-th of the values that they should be (from ordering the combined test statistics), and three-way Fisher Integration yields values that are 1/10,000-th the values that they should be (Fig. 4A). By contrast, when using the Brown method, r ≈ 1 in all cases (Fig. 4B).

**Figure 4:**
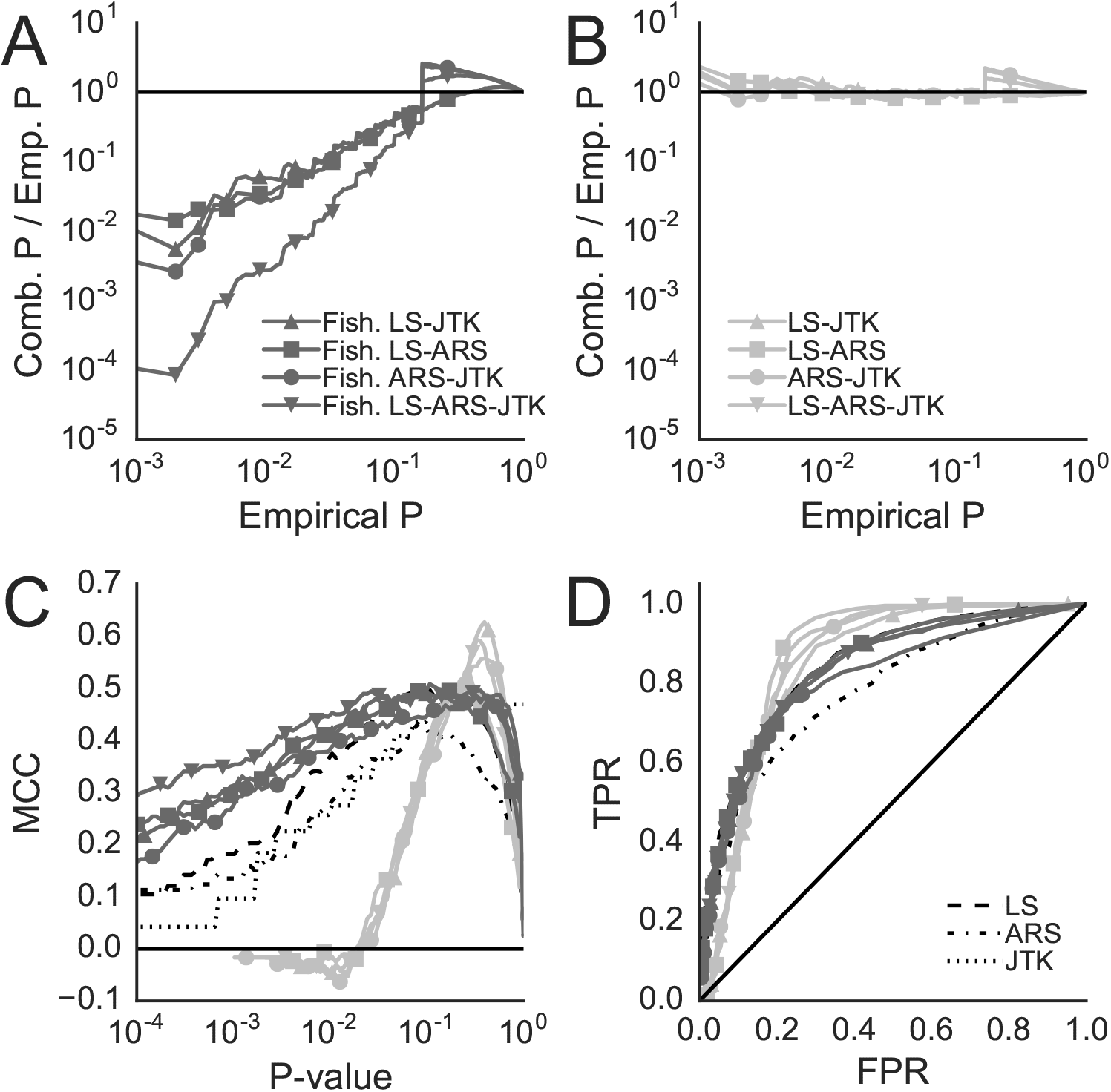
Combining methods does not improve rhythm detection. (A and B) Indicated rhythm detection methods were tested on 1000 Gaussian noise time series; the horizontal black line is *r* = 1. (A) P-values from Fisher Integration of Lomb-Scargle (LS), ARSER (ARS), and JTK_CYCLE (JTK) after adjusting their p-values to be uniform (Fig. 2). (B) P-values from the Brown integration of the same methods with adjusted p-values in (A). (C and D) To test the performance of the indicated rhythm detection methods, we generated rhythmic time series as described in the text and combined them with the Gaussian noise time series. (C) Matthews Correlation Coefficient (MCC) for classification of the randomly generated time series as arrhythmic and the cosine-generated time series as rhythmic for different p-value thresholds. The p-values for the Fisher-combined methods are not true p-values, as seen in (A); they are shown here to reflect that the Fisher-combined methods appear to do well if their p-values are not realized to be incorrect. (D) Receiver operating characteristic: comparison of True Positive Rate (TPR) to False Positive Rate (FPR) as significance threshold is varied. The horizontal line in (C) and the diagonal line in (D) indicate random classification.

Having corrected the p-value integration, we compared the combinations of the methods to the individual methods for rhythm detection. We generated 396 rhythmic time series by adding Gaussian noise plus 10 (to avoid negative values) to a cosine with 24 points sampled evenly across 2 periods (analogous to sampling every 2 hours for 48 hours across 2 24-hour periods); the standard deviation of the Gaussian noise was equal to the amplitude of the cosine. For these time series, the *R*^2^ between empirically-corrected p-values produced by JTK_CYCLE and Lomb-Scargle is *R*^2^ = 0.468, that between JTK_CYCLE and ARSER is *R*^2^ = 0.297, and that between Lomb-Scargle and ARSER is *R*^2^ = 0.431. To analyze the classification strength of the different methods, we combined the randomly generated data with the cosine-generated data, yielding a dataset with ≈ 28% rhythmic time series, which is in line with tissue-specific percent rhythmicity estimates [7]. While the combinations of the methods integrated improperly with the Fisher method appear to outperform the individual methods at p-values below typical significance cutoffs (Figs. 4C and D, dark gray), once the p-values are accurately calculated using the Brown method, the combined methods (Figs. 4C and D, light gray) underperform the individual methods for low p-values. In Fig. 4C this is shown using the Matthews Correlation Coefficient (MCC), which is a scalar metric that summarizes a confusion table [10] where 1 indicates a perfect classifier and 0 indicates a classifier that does no better than random. While the scores for the Fisher-integrated combinations are above those of the individual methods, the scores for the Brown-integrated combinations are below those of the individual methods for *p* < 0.1. These trends can also be seen by plotting the True Positive Rate (TPR) as a function of the False Positive Rate (FPR) as the significance threshold is varied (the dark gray curves in Fig. 4D are above the light gray curves for *p* < 0.1).

## 3 Conclusions

During rhythm detection, it is important to account for correlations in p-values that arise from the application of multiple tests to the same experimental time series. Here, we show that this issue impacts the correction for testing across reference waveforms in JTK_CYCLE [5] and RAIN [13], and it impacts the integration of p-values from multiple methods in MetaCycle [14]. While it can be computationally costly to correct the p-values empirically [6, 7], Brown's correction of Fisher integration involves little additional effort [2]. Surprisingly, we find no advantage to combining methods with accurate p-values. This may be due to the fact that most rhythm detection methods overlap in the aspects of the time series that they examine. This high level of correlation results in a type of multiple hypothesis comparison problem that increases the p-values without likewise increasing the sensitivity of the combined methods to detect rhythmicity. For method combination to be effective, new methods of rhythm detection would need to be developed that approach rhythm detection from new directions.

## Acknowledgments

This work was completed in part with resources provided by the University of Chicago Research Computing Center.

## References

[1] Yoav Benjamini and Yosef Hochberg. Controlling the False Discovery Rate: A Practical and Powerful Approach to Multiple Testing. Journal of the Royal Statistical Society, Series B (Methodological), 57(1):289–300, 1995.

[2] Morton B Brown. A Method for Combining Non-Independent, One-Sided Tests of Significance. Biometrics, 31(4):987–992, 1975.

[3] Anastasia Deckard, Ron C Anafi, John B Hogenesch, Steven B Haase, and John Harer. Design and analysis of large-scale biological rhythm studies: a comparison of algorithms for detecting periodic signals in biological data. Bioinformatics, 29(24):3174–80, Dec 2013.

[4] R A Fisher. Statistical Methods for Research Workers. Oliver and Boyd, 1925.

[5] Michael E Hughes, John B Hogenesch, and Karl Kornacker. JTK_CYCLE: an efficient nonparametric algorithm for detecting rhythmic components in genome-scale data sets. Journal of Biological Rhythms, 25(5):372–80, Oct 2010.

[6] Alan L. Hutchison, Mark Maienschein-Cline, Andrew H. Chiang, S. M Ali Tabei, Herman Gudjonson, Neil Bahroos, Ravi Allada, and Aaron R. Dinner. Improved Statistical Methods Enable Greater Sensitivity in Rhythm Detection for Genome-Wide Data. PLoS Comput Biol, 11(3):e1004094, 2015.

[7] Alan Louis Hutchison, Ravi Allada, and Aaron R. Dinner. Bootstrapping and Empirical Bayes Methods Improve Rhythm Detection in Sparsely Sampled Data. In preperation, 2017.

[8] Kevin P. Keegan, Suraj Pradhan, Ji Ping Wang, and Ravi Allada. Meta-analysis of Drosophila circadian microarray studies identifies a novel set of rhythmically expressed genes. PLoS Computational Biology, 3(11):2087–2110, 2007.

[9] N R Lomb. Least-squares frequency analysis of unequally spaced data. Astrophysics and Space Science, 39(2):447–462, 1976.

[10] B.W. Matthews. Comparison of the predicted and observed secondary structure of T4 phage lysozyme. Biochimica et Biophysica Acta (BBA) - Protein Structure, 405(2):442–451, Oct 1975.

[11] J. D Scargle. Studies in astronomical time series analysis. II - Statistical aspects of spectral analysis of unevenly spaced data. APJ, 263(Dec):835–853, 1982.

[12] John D. Storey. The Positive False Discovery Rate: A Bayesian Interpretation and the q-Value. The Annals of Statistics, 31(6):2013–2035, 2003.

[13] Paul F Thaben and Pal O Westermark. Detecting rhythms in time series with RAIN. Journal of Biological Rhythms, 29(6):391–400, 2014.

[14] Gang Wu, Ron C Anafi, Michael E Hughes, Karl Kornacker, and John B Hogenesch. MetaCycle: an integrated R package to evaluate periodicity in large scale data. Bioinformatics, 1-3(July):040345, 2016.

[15] Gang Wu, Jiang Zhu, Jun Yu, Lan Zhou, Jianhua Z Huang, and Zhang Zhang. Evaluation of five methods for genome-wide circadian gene identification. Journal of Biological Rhythms, 29(4):231–42, 2014.

[16] Rendong Yang and Zhen Su. Analyzing circadian expression data by harmonic regression based on autoregressive spectral estimation. Bioinformatics, 26(12), 2010.

